# AVITI sequencing of a four-generation CEPH/Utah pedigree confirms low mutation rates at homopolymer loci despite their low sequence complexity

**DOI:** 10.1101/2025.09.25.678675

**Authors:** Hannah C. Happ, Thomas A. Sasani, Derek Warner, Deborah W. Neklason, Aaron R. Quinlan

**Author notes:** Correspondence: Hannah C. Happ, Aaron R. Quinlan.

## Abstract

**Background:** Short tandem repeats (STRs) and homopolymers are among the most mutable loci in the human genome. Despite their presumed mutability owing to replication slippage, homopolymer loci exhibit lower mutation rates and minimal paternal age effects compared to other STRs. This paradox questions if technical limitations, rather than biological mechanisms, explain these observations.

**Results:** We used the Element Biosciences AVITI platform to sequence the genomes of a 48-member, four-generation CEPH/Utah pedigree. As the AVITI platform reduces error rates at repetitive sequences compared to Illumina, this design enabled accurate mutation discovery at 90% of assayed homopolymers and a 1.7-fold increase in discoverable mutations compared to Illumina. We identified a median of 35 *de novo* homopolymer mutations per trio and a mutation rate of 5.28 × 10^−5^ DNMs per locus per generation, confirming a lower rate than dinucleotides (1.94 × 10^−4^). Most DNMs were single base-pair expansions or contractions. Despite comprising <1% of homopolymer loci, G/C homopolymers showed 18-fold higher mutation rates than A/T homopolymers; in contrast, the high dinucleotide mutation rate is not driven by a particular motif class. Parent-of-origin analysis revealed 78% of homopolymer mutations are paternal in origin, but no significant paternal age effect was observed.

**Conclusions:** This study confirms that homopolymers exhibit lower mutation rates and lack strong paternal age effects compared to other STRs, likely owing to the combination of a lower propensity to form slippage-causing secondary structures and more efficient mismatch repair. Our set of high-quality mutations suggest these phenomena are biological rather than technical in nature. Finally, we demonstrate that AVITI sequencing unlocks previously intractable regions of the genome and will be a powerful tool for continued investigation of repeat mutation.

## Background

*De novo* mutations (DNMs) are the fundamental source of genetic diversity, yet they are also the cause of nearly 60% of genetic disorders and underlie many other genetic diseases (1–4). By sequencing the genomes of many thousands of families, many of the factors that influence single-nucleotide (1,5–10) and structural (11–13) mutation are now well understood. However, our understanding of some of the most mutable loci in the human genome, short tandem repeats (STRs), remains comparatively limited. STRs comprise 1-6 base pair (bp) motifs repeated sequentially, and they account for approximately 3% of the human genome (14,15). Despite their relatively small genomic footprint, STRs contribute substantial genomic diversity due to their extremely high mutation rates, which are orders of magnitude higher than those observed at high complexity regions of the genome (16,17). This high mutability is largely understood to be due to high rates of polymerase slippage during DNA replication (18,19). During replication, repetitive single-stranded DNA has a high propensity to form secondary structures such as hairpins, cruciforms, and Z-DNA (20). These bulky, non-B DNA structures cause the polymerase to stall and release from the template strand; reassembly frequently results in misalignment of the template and nascent strand, which can lead to an addition or deletion of one or more repeat units (21–23). Unfortunately, this same mechanism plagues polymerase-based sequencing methods. Polymerase slippage during DNA synthesis involved in library preparation and sequencing creates a distribution of allele sizes that differ by multiples of the repeat unit (24,25); this “stutter noise” can obscure the underlying genotype, making it difficult to distinguish true DNMs from technical artifacts. This is especially true for homopolymers (26) which are frequently excluded or analyzed with caution. This exclusion has limited our understanding of mutation dynamics at these loci.

In 2023, Element Biosciences introduced the AVITI sequencing platform, which achieves lower error rates in and around repetitive sequences compared to existing technologies (27,28). Rather than sequencing by synthesis, the AVITI uses sequencing by “avidity base chemistry”, which separates polymerase procession from base identification. After a nucleotide is added to the growing DNA chain, fluorescently labeled “avidites” bind the DNA template with high specificity and stability; as a result, dissociation during imaging is reduced, enabling more accurate base calling. The AVITI platform also uses rolling circle amplification, which reduces error propagation. Together, these features make AVITI a compelling platform for studying STR mutation. Therefore, we sought to use the Element AVITI to investigate the dynamics of *de novo* mutation at STRs, with a focus on homopolymers. To achieve this goal, we used AVITI to sequence the genomes of 48 members of a four-generation CEPH/Utah pedigree, which provides a total of 39 meioses for DNM discovery (**Figure 1**). Sixteen members of the pedigree were also sequenced on an Illumina platform as part of the Platinum Pedigree consortium (29).

**Figure 1.**
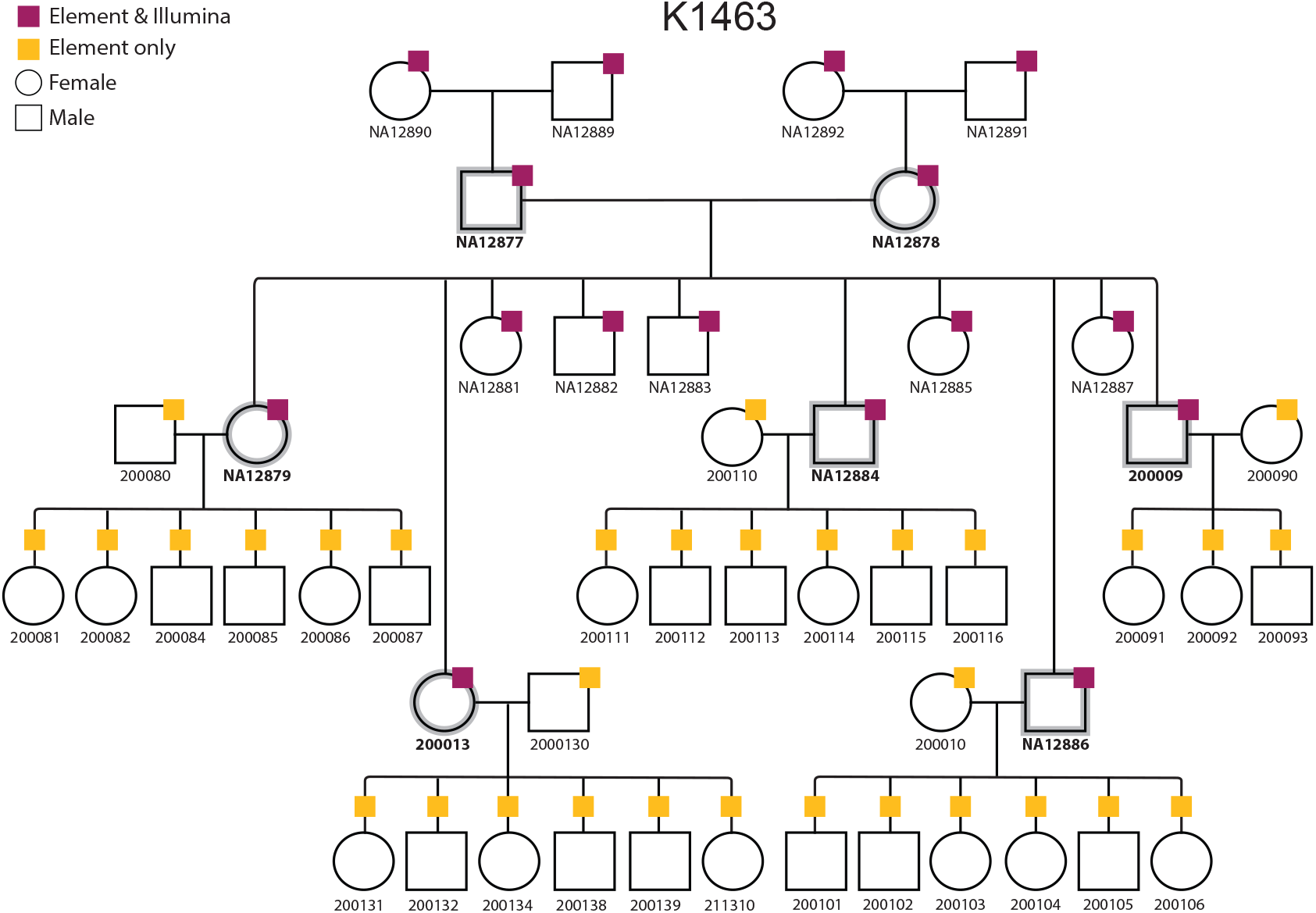
CEPH K1463 Platinum Plus pedigree. We generated whole genome sequencing (WGS) data for 48 members of a 4-generation CEPH/Utah pedigree with Element Biosciences’ AVITI platform (yellow). For a subset of the cohort (n=16), we also generated Illumina WGS data (pink). For seven individuals (gray highlight) whose parents and children are sequenced, DNMs can be assigned to a parent of origin.

By combining DNA from a large, multi-generational pedigree with a DNA sequencing technology that minimizes sequencing errors at repetitive loci, this study design enables accurate genome-wide estimates of homopolymer DNM counts and rates. Moreover, the resulting dataset empowers the investigation of two curious patterns of homopolymer mutation that have been reported in previous studies. First, prior studies observe that STR mutation rates generally decrease as motif length increases (30,31). Following this pattern, one might expect homopolymers to have the highest mutation rate of STRs; however, previous WGS-based studies report the highest STR mutation rate at dinucleotide repeats (29,32–34). Additionally, previous studies observe a paternal age effect at nonhomopolymer STRs, in which older fathers transmit more DNMs than younger fathers (32,34,35). In contrast, no or an extremely low paternal age effect has been detected at homopolymers (34,35). This observation is puzzling since STR mutations occur during replication, and spermatogonial stem cells replicate and divide continuously beginning in puberty; thus, replication-associated errors, including slippage events, are predicted to increase with paternal age. We have the unique opportunity to reliably test whether previous estimates of homopolymer mutation rates and how they vary as a function of paternal age are biologically-sound or the result of technical limitations.

## Results

### Element sequencing provides better substrate for identifying DNMs at homopolymers

We sequenced the genomes of 48 members of the CEPH K1463 pedigree with the Element Bioscience’s AVITI platform to a genome-wide median sequencing depth of 46X; a subset of 16 members of the pedigree were also sequenced with the Illumina platform to a median depth of 36X (**Supplementary Table 1**). On a per-trio basis, we used HipSTR-generated genotypes to query 777,273 autosomal homopolymer loci for DNMs, conditional upon sufficient genotype quality and sequencing depth for all three members of the trio (**Methods**). Overall, the Element technology led to HipSTR genotype predictions for each member of a mother, father, and child trio and testing for a DNM at up to 90.1% of homopolymer loci (median: 89.4%, range: 62.1 - 90.1%) in our reference set (**Figure 2**). This represents a 1.7-fold increase in discoverable DNMs compared to trios sequenced on an Illumina platform (**Figure 2**). The lower cluster of trios in the Element cohort is driven by one G3 parent with a low median insert size (**Supplementary Table 1**), which impacts the number of loci at which HipSTR can accurately genotype.

**Figure 2.**
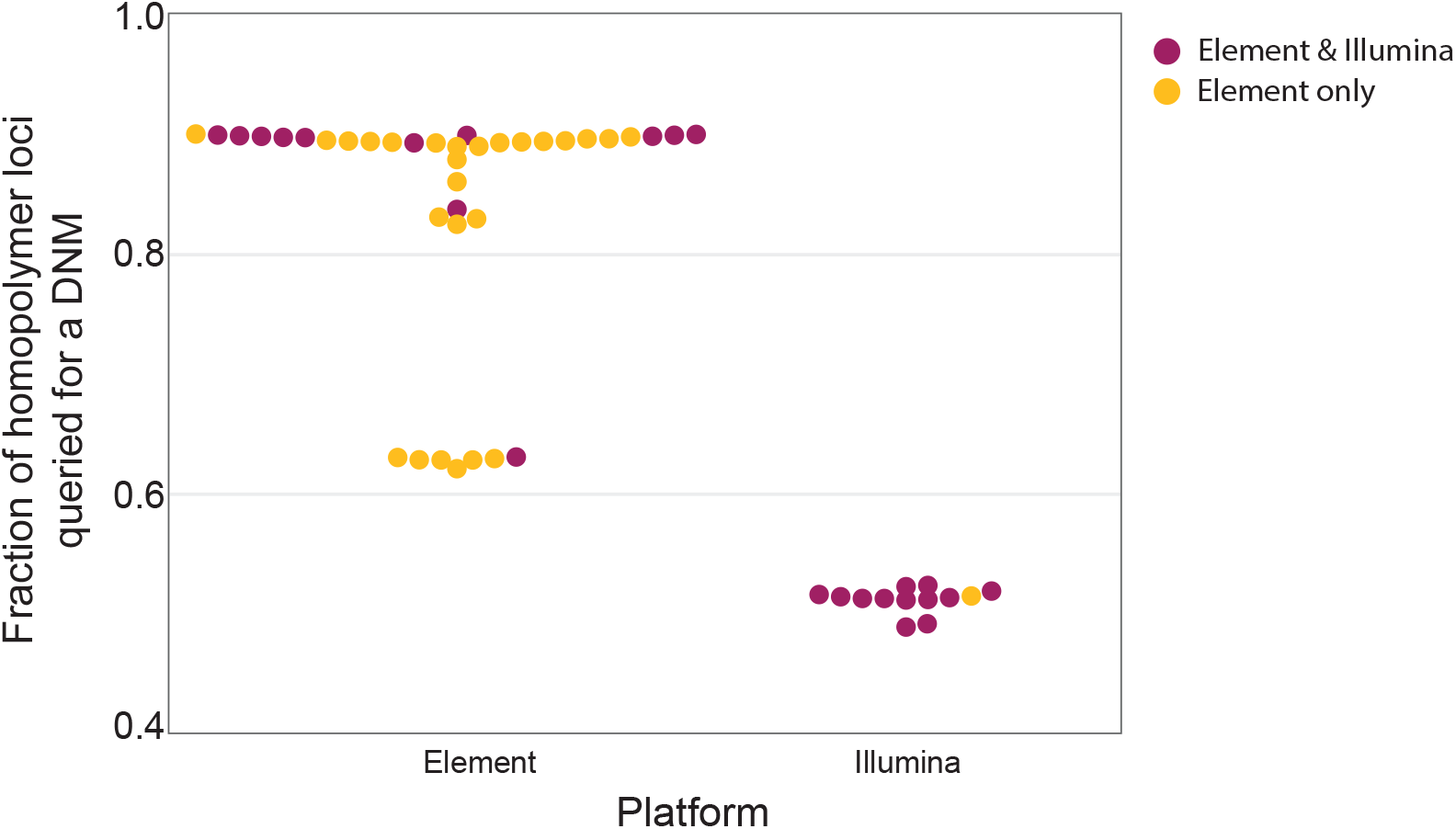
Fraction of homopolymer loci that could be queried for a DNM in each trio, stratified by sequencing platform. We queried 777,273 autosomal homopolymer loci for DNMs across 39 trios. The fraction of loci at which a DNM is discoverable is plotted as a function of sequencing technology. Trios were sequenced with Element only (yellow) or both Element and Illumina platforms (pink).

### Using high-confidence homopolymer DNMs to characterize mutation dynamics

Across 39 meioses, we observe a median of 35 homopolymer DNMs per trio (range: 15–59) and estimate a mutation rate of 5.28 × 10^−5^ per locus per generation (95% CI = 5.01 × 10^−5^ - 5.57 × 10^−5^) (**Figure 3**). Extending these analyses to nonhomopolymer STRs, we observe a median of 59 nonhomopolymer DNMs per trio (range: 17-109) and estimate a mutation rate of 9.57 × 10^−5^ per locus per generation (95% CI = 9.70 × 10^−5^ - 1.05 × 10^−^4). Moreover, when we look across all STRs, we observe the highest mutation rate at dinucleotide repeats at 1.77 × 10^−4^ (95% CI = 1.85 × 10^−^4 - 2.04 × 10^−4^) (**Table 1)**. These results align with several previous reports of STR mutation rates (29,32–34) (**Supplementary Figure 1**).

**Figure 3.**
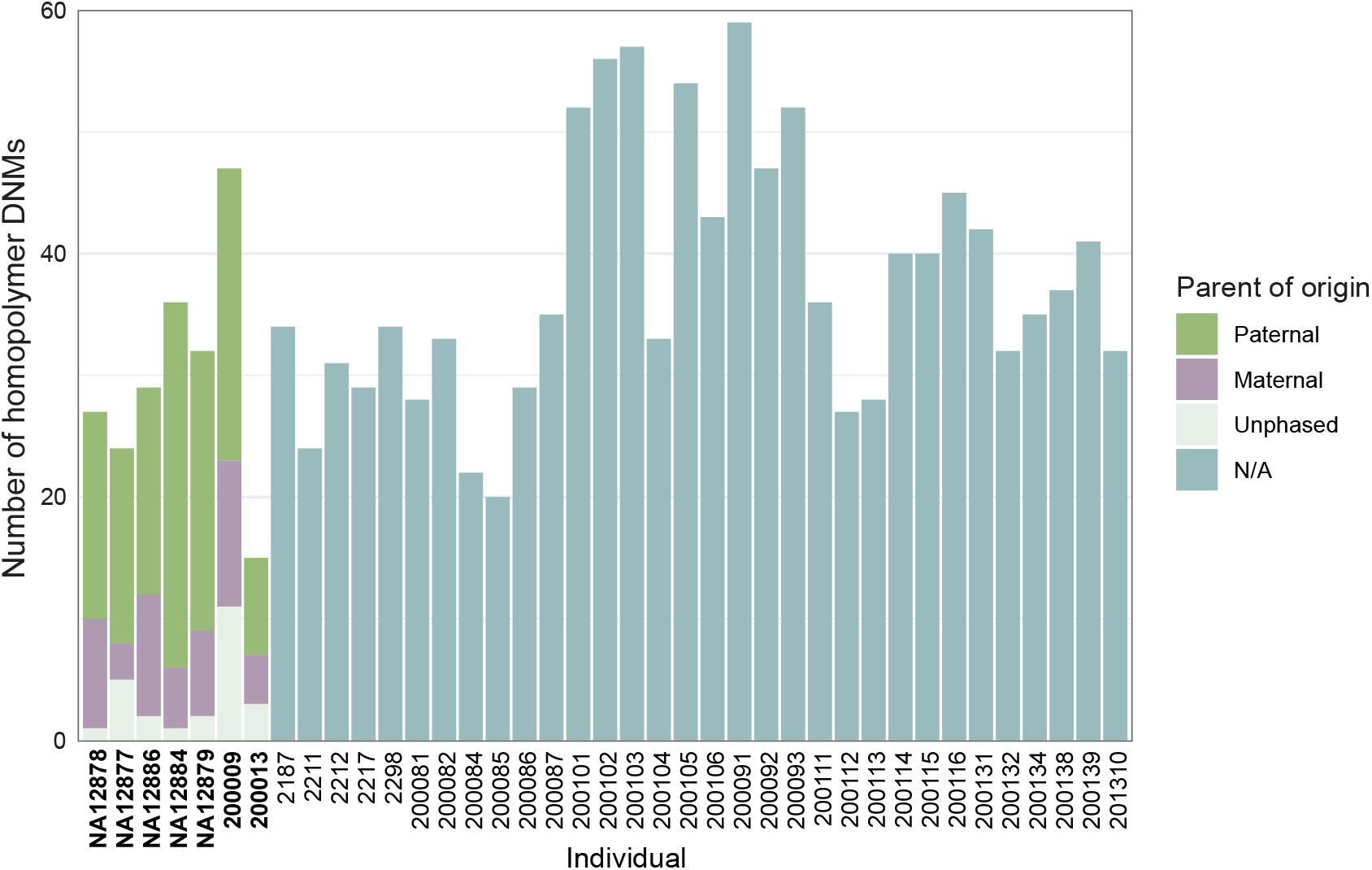
Homopolymer DNMs counts. Bars represent the number of homopolymer DNMs identified per individual. For seven individuals (bold) whose parents and children were also sequenced, DNMs could be assigned to a parent of origin using haplotype tracing.

**Table 1.** Mutation rates across all STR motif sizes.

| Motif size (bp) | DNMs | Queried sites | Mutation rate (per locus per generation) | 95% CI |
| --- | --- | --- | --- | --- |
| 1 | 1417 | 26814603 | $5.28 \times 10^{-5}$ | $5.01 \times 10^{-5} - 5.57 \times 10^{-5}$ |
| 2 | 1644 | 8469259 | $1.94 \times 10^{-4}$ | $1.85 \times 10^{-4} - 2.04 \times 10^{-4}$ |
| 3 | 95 | 2358520 | $4.03 \times 10^{-5}$ | $3.26 \times 10^{-5} - 4.92 \times 10^{-5}$ |
| 4 | 475 | 7024512 | $6.76 \times 10^{-5}$ | $6.17 \times 10^{-5} - 7.40 \times 10^{-5}$ |
| 5 | 73 | 2974032 | $2.45 \times 10^{-5}$ | $1.92 \times 10^{-5} - 3.09 \times 10^{-5}$ |
| 6 | 33 | 2135289 | $1.55 \times 10^{-5}$ | $1.06 \times 10^{-5} - 2.17 \times 10^{-5}$ |
| All STRs | 3737 | 49776215 | $7.51 \times 10^{-5}$ | $7.27 \times 10^{-5} - 7.75 \times 10^{-5}$ |
| Nonhomopolymers | 2320 | 22961612 | $1.01 \times 10^{-4}$ | $9.70 \times 10^{-5} - 1.05 \times 10^{-4}$ |

Moreover, this set of DNMs enables us to investigate homopolymer DNM rates in the context of allele length and nucleotide content. The analyses can also help tease apart the biological underpinnings of homopolymer mutation dynamics.

First, to determine whether homopolymer mutation rates varied as a function of length, we compared DNM counts across bins of reference allele lengths using a Poisson regression model, adjusting for the number of queried sites in each bin. Since we observe a strong correlation between the reference allele length and parental allele lengths (Pearson *r* = 0.88; **Supplementary Figure 2**), we used the reference allele length as a proxy for the length of the allele that mutated. In general, mutation rates increase with longer reference alleles, with the exception of one bin (70-84 bp) (**Supplementary Table 2**). The shortest bin (10-24 bp) had a mutation rate of 3.12 × 10^−5^ per site (95% CI: 2.90 × 10^−5^ - 3.36 × 10^−5^). The longest bin (85-99 bp) had a mutation rate of 3.65 × 10^−3^ (95% CI: 7.53 × 10^−4^ - 1.07 × 10^−2^), which is a 116-fold increase in mutation rate compared to that of the alleles in the shortest bin.

Additionally, we observe striking effects of nucleotide composition on homopolymer mutation rates. Although G/C homopolymers make less than <1% of all homopolymer loci, these sites are enriched for DNMs; we observe an 18-fold higher mutation rate compared to A/T homopolymers (rate ratio = 0.056; 95% CI: 0.042-0.077; *P* < 2.2 × 10^-16^) (**Figure 4**). Given the strong motif-specific effect we observe on G/C homopolymer mutation rates, we asked whether the mutation rates of particular dinucleotide motifs are driving the high overall dinucleotide mutation rate. This analysis revealed two interesting results. First, we find that CG-dinucleotides are depleted among all dinucleotide motifs, which is consistent with their overall depletion in the genome due to spontaneous deamination of methylated cytosines at CpG dinucleotides. On average, only 115 CG/GC dinucleotides per trio were queried for a DNM; we did not observe any DNMs at these loci. Second, we find that all other dinucleotide motif classes have overlapping or significantly higher mutation rates compared to homopolymers, suggesting the high dinucleotide mutation rate is not driven by particular motif-specific effects (**Figure 4**). Finally, we observe similar mutation rates at homopolymers and trinucleotide repeats, which are similarly not driven by motif-specific rates (**Supplementary Figure 3**).

**Figure 4.**
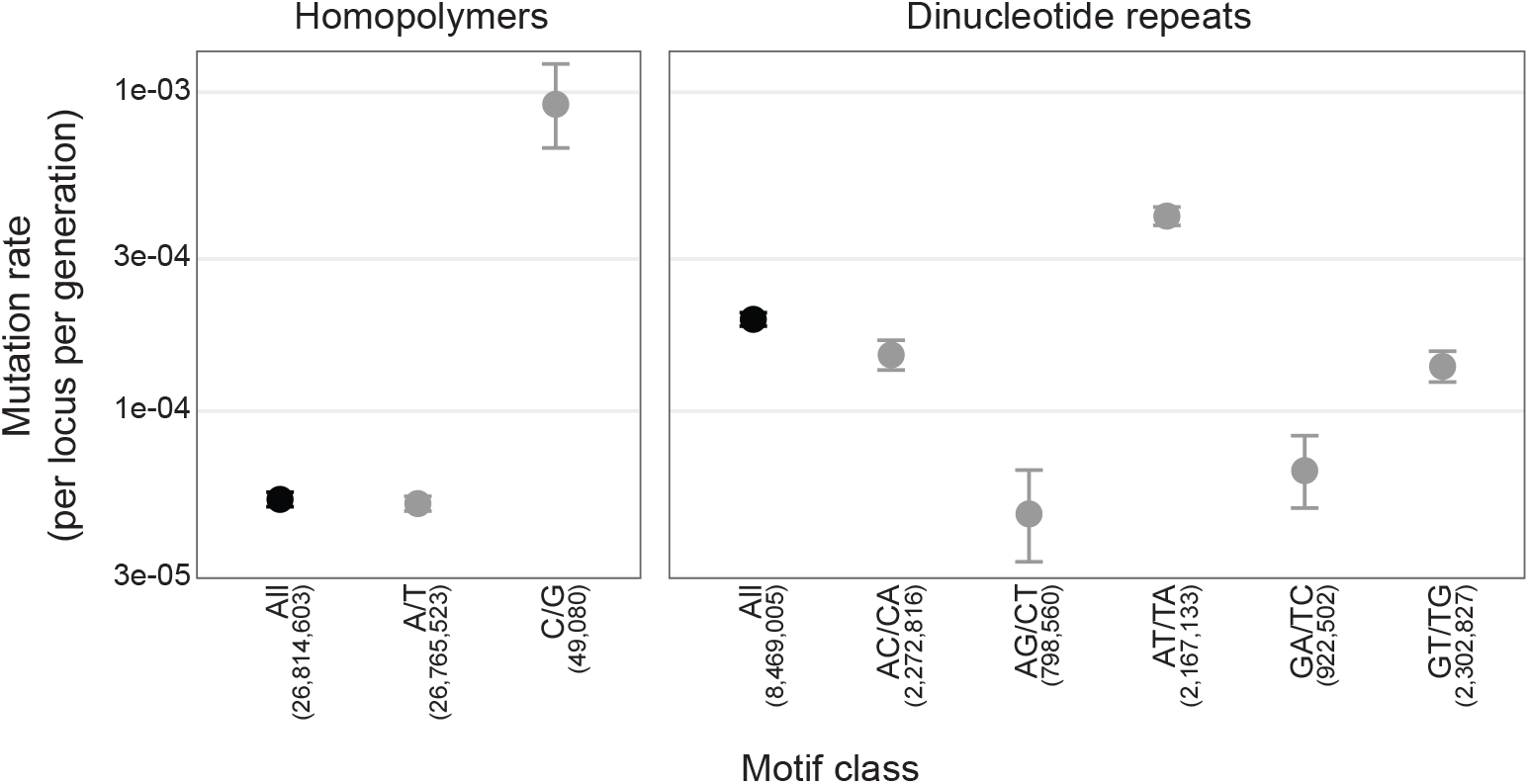
Homopolymer and dinucleotide repeat mutation rates as a function of motif class. Homopolymers were grouped by base content (A/T, G/C), dinucleotide motifs were collapsed by reverse complement (e.g. AG = CT). Motif classes for which no DNMs were observed are not shown. Error bars represent 95% confidence intervals. The number of loci queried for a DNM across 39 meioses (denominator) for each motif class is shown beneath each motif class.

### Phasing of DNMs in a subset of family members reveals no parental age effects on homopolymer mutation rates

For the seven individuals in the pedigree for whom we sequenced both parents and offspring (**Figure 1**), we used transmission to the third generation to phase validated DNMs to a parent of origin and to a particular parental haplotype. We determined the parental haplotype of origin for 91% (range: 83-99%) of DNMs in these individuals. From this set of DNMs, we observe a ratio of paternal to maternal DNMs of 3.37:1 and find that, like SNV mutations, 78% of DNMs are paternal in origin (**Figure 3**). These data further enable us to interrogate the effect of parental age on homopolymer DNM rates. After fitting Poisson regressions, we observe no statistically significant effect of paternal or maternal age on homopolymer DNM rates (**Figure 5A**) (*P* = 0.143 and 0.125 for paternal and maternal age, respectively). Consistent with previous findings, we do observe a significant association between paternal age and nonhomopolymer DNM rates (*P* = 0.0256); however, we are underpowered to detect a significant maternal age effect on nonhomopolymer DNM rates (*P* = 0.786).

**Figure 5.**
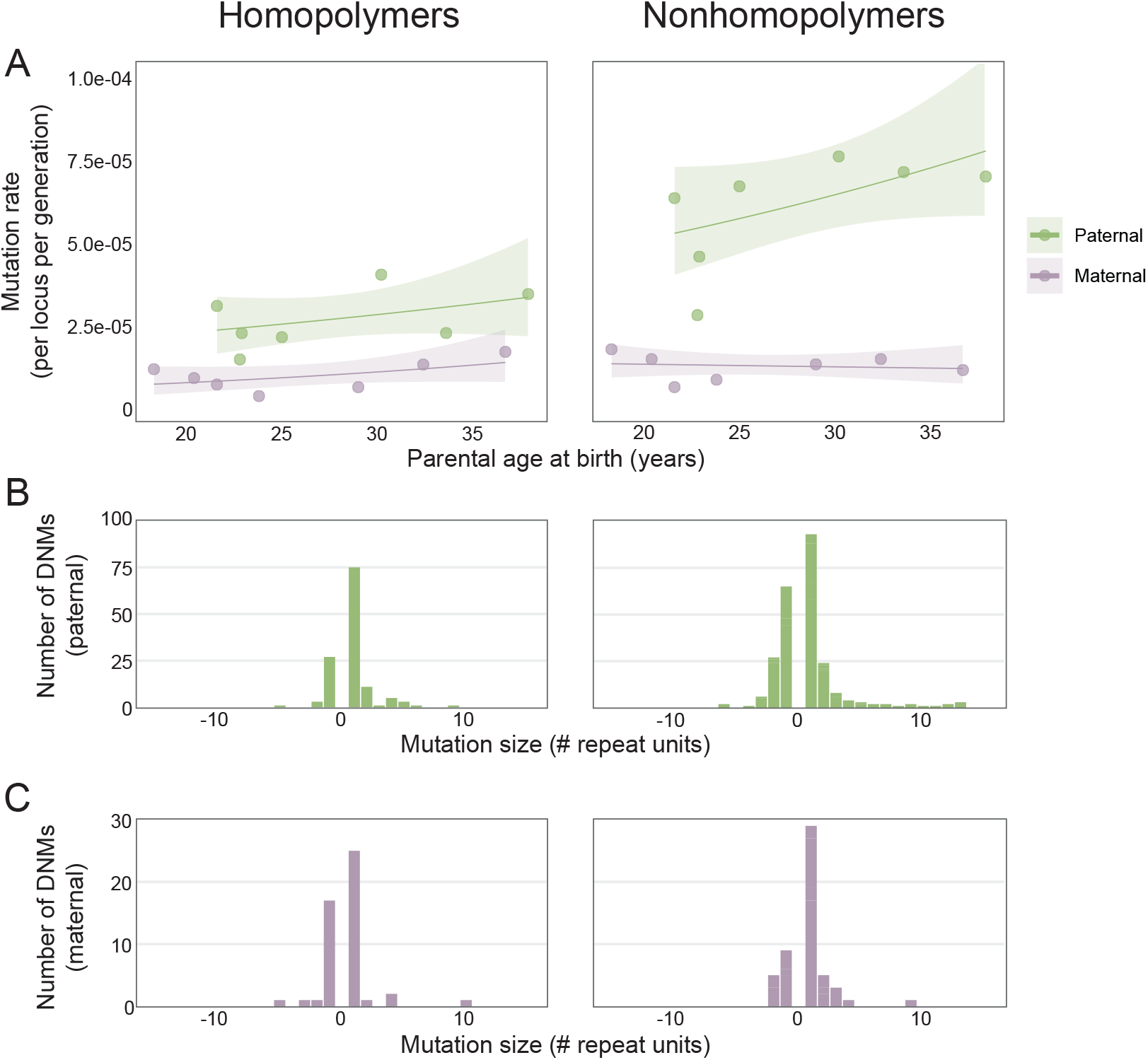
Distribution of parental age effects and DNM mutation size. **A)** Homopolymer (motif size = 1 bp) and nonhomopolymer (motif size = 2-6bp) mutation rates as a function of parental age at birth. Poisson regressions were fit for mothers and fathers separately using a log link (with 95% confidence bands). **B**,**C)** Homopolymer and nonhomopolymer DNMs that could be assigned at a parental haplotype of origin were used to ascertain the distribution of expansion and contraction sizes.

We further leveraged these phased DNMs to interrogate the size distributions of homopolymer mutations. We observed that among both maternal- and paternal-origin DNMs, most mutations are 1 bp expansions or contractions, with a bias toward the former (two-sided binomial test, *P* = 1.4 × 10^−8^) (**Figure 5B,C**). Additionally, nonhomopolymers similarly exhibit a bias toward expansions (two-sided binomial test, *P* = 2.597 × 10^−5^), most commonly single repeat unit expansions (**Figure 5B,C**).

## Discussion

Homopolymers are frequently excluded from germline mutation studies or treated with caution due to high error rates observed at these loci. Leveraging Element Bioscience’s AVITI platform’s improved accuracy at repetitive regions and a large, four-generation pedigree, we sought to robustly evaluate mutation dynamics at homopolymers. Our findings, particularly the low mutation rate at homopolymers compared to dinucleotide repeats, as well as the absence of a strong paternal age effect on mutation rate, suggest a biological, rather than technical, signal.

### A low homopolymer mutation rate or a high dinucleotide mutation rate?

Consistent with previous sequencing-based studies of STR mutation, we observe the highest mutation rates at dinucleotide repeats, and lowest mutation rates at hexanucleotide repeats (**Supplementary Figure 1**) (29,32–34,36). Following this pattern, one might expect homopolymers to have the highest mutation rates; however, our results confirm prior evidence to the contrary. The high mutation rate at dinucleotides compared to homopolymers is likely in part due to the types of secondary structures these repeats are prone to form during replication (37). Dinucleotides, in particular inverted repeats such as TA-tracts, are more likely to form hairpins or cruciforms than homopolymers (37,38). Consistent with this notion, we observe the highest mutation rate at TA-dinucleotides. Additionally, dinucleotides, unlike homopolymers, can form Z-DNA structures, which require alternating purine-pyrimidine nucleotides (e.g. CA/TG, AT/TA). Importantly, homopolymers are not immune to secondary structures; poly(G)-tracts can form stable G-quadruplexes, which may underlie the significant enrichment of DNMs that we observe at G/C homopolymers compared to A/T tracts (23). Considering our results in the context of the high propensity of dinucleotide repeats to form secondary non-B DNA structures reframes what we originally considered a curious observation; the low homopolymer mutation rate likely reflects a lower propensity of these tracts to form slippage-causing secondary structures compared to dinucleotide tracts.

In addition to differences in secondary structure formation, the discrepancy in mutation rate between homopolymers and dinucleotides is also likely due, in part, to differential damage repair. Replication slippage errors at STRs are normally surveilled and corrected by DNA mismatch repair (MMR) mechanisms (39). Defects in MMR proteins in cells and tissues lead to microsatellite instability (40–42) such that under normal conditions most slippage events at STRs are efficiently repaired before they can become heritable mutations. Components of the MMR machinery have differential specificity for DNA lesions, which plays a role in shaping the varying mutation landscape across STRs. For example, in eukaryotes two MutS complexes initiate MMR at different types of errors: MutSα (MSH2-MSH6) primarily targets single-base mismatches and 1-bp loops, and MutSβ (MSH2-MSH3) primarily repairs loops of two or more bases but can also repair 1-bp loops (43–45). Furthermore, prior studies of mutation spectra in the absence of MMR report a mutational bias toward deletions, which suggests under wild-type conditions, deletions are more readily repaired than insertions at homopolymers (46–49). Our results are consistent with this notion, as we observe more insertions than deletions among all STRs. Additionally, the relatively high abundance of homopolymers compared to other STRs in conjunction with the high frequency of 1-bp indels, suggests that 1-bp indels at homopolymers likely account for the most common mismatch generated at these loci. The redundancy of MutSα and MutSβ in recognizing 1-bp indels may ensure these mutations are readily identified and repaired and may account, in part, for the lower mutation rate at homopolymers compared to dinucleotide repeats.

Ultimately, our observation of a lower mutation rate at homopolymers compared to dinucleotide repeats should be considered in the context of both secondary structure formation and mismatch repair. Dinucleotides are especially prone to forming bulky, non-B DNA structures that create opportunities for replication errors. Moreover, MMR pathways redundantly target 1-bp indels, the most frequent homopolymer mutation, correcting these errors before they can be propagated as DNMs. Together, the higher structural instability of dinucleotides and the enhanced repair of homopolymer errors may explain why dinucleotide mutation rates exceed those of homopolymers.

### An unexpected lack of paternal age effect on homopolymer DNMs

Given our ability to phase DNMs to a parent of origin using transmission to a subsequent generation, we were eager to interrogate parental age effects on homopolymer mutation rates. Consistent with findings from studies of SNVs (5,7–10) and nonhomopolymer STRs (32–35), we observe a paternal-to-maternal ratio of 3.37:1, with 78% of homopolymer DNMs arising on the paternal haplotype. Surprisingly, however, we observe no significant effect of paternal age on the homopolymer DNM rate. Given that spermatogonial stem cells continuously divide throughout a man’s reproductive years and homopolymer mutation is thought to be driven by polymerase slippage during replication, we might expect mutation rates to increase with paternal age. Two recent studies specifically interrogate parental age effects on homopolymer DNM rates (34,35). One reported no paternal age effect, and the other, powered by a substantially larger dataset than former studies, observed a weak but significant paternal age effect. One conclusion might be that this study is underpowered to detect a paternal age effect on homopolymer DNM rates, however, we and other studies do detect a significant paternal age effect on nonhomopolymer STR DNM rates (32–35). Prior studies have posited that the lack of a paternal age effect on the former may reflect difficulties in accurately genotyping homopolymers and disentangling age effect signal from sequencing noise. Given the increased accuracy of our data, the attenuated paternal age effect on the former is particularly curious and likely representative of underlying biological processes. For example, homopolymer DNMs may occur during windows of germ-cell development that are age-independent or be repaired more efficiently than nonhomopolymers during proliferative stages due to differences in expression or saturation of various repair pathways. Moreover, poly(A/T)-tracts are associated with low nucleosome-occupancy, which may lead to increased exposure to both DNA damage and surveillance/repair machinery (50). Additional work to unravel this curiosity is warranted.

## Conclusions

Our study offers an opportunity to confidently measure homopolymer mutation dynamics, which are typically reported with caution. These results are powered by a substantial improvement in the ability to discover homopolymer DNMs using AVITI sequencing. Illumina-based sequencing is plagued by stutter artifacts and poor alignment at homopolymers; as a result, STR variant detection tools like HipSTR cannot accurately genotype many of these alleles, and a DNM at such an allele would go undiscovered. In contrast, less stutter noise in the Element sequencing enables high-quality genotyping in all three trio members of the vast majority of the homopolymers in our reference set. As the AVITI platform is new, it is not yet as widely available as other platforms. However, our study demonstrates that it is an ideal choice for studies of, for example, microsatellite instability and the genetic basis of neurological disorders, where characterizing STR and homopolymer mutations is imperative. An important caveat, however, is that AVITI read lengths are too short to accurately characterize mutations in very long STRs. The high-quality catalog of homopolymer DNMs will be a valuable benchmark dataset for development of new and refinement of current STR genotyping tools. These tools, coupled with the Element AVITI platform, will be a powerful combination for future studies of homopolymers and other highly complicated but low-complexity genomic elements.

## Methods

### Cohort and sequencing

Informed consent was obtained from the CEPH/Utah participants. The University of Utah Institutional Review Board approved the study (University of Utah IRB protocol IRB_00065564). 28 family members were re-engaged or newly enrolled as previously described (29). This study includes an extended version of the previously described pedigree. The additional family members were similarly re-engaged (n = 2) to update informed consent and health history or newly enrolled (n = 18). All individuals are broadly consented for scientific purposes.

Archived DNA from individuals in G1-3 was extracted from whole blood. For newly enrolled family members, DNA was extracted from whole blood using the Flexigene system (Qiagen, Cat# 51206). Element genome sequencing data (Element) were generated according to the manufacturer’s recommendations. In brief, PCR-free libraries were prepared using mechanical shearing (Covaris, Cat# 500295), yielding ∼350 bp fragments, and the Element Elevate library preparation kit (Element Biosciences, Cat# 830-00008). Linear libraries were quantified by quantitative PCR and sequenced with either the Cloudbreak Freestyle (*n* = 20, Element Biosciences, Cat#) or Cloudbreak UltraQ (*n* = 28, Element Biosciences, Cat#) sequencing kits, which can achieve Q40 or Q50 quality data, respectively. Bases2Fastq Software (Element Biosciences) was used to generate demultiplexed FASTQ files. Demultiplexed FASTQ files were aligned to a GRCh38 reference using BWA (51). Illumina genome sequencing data (Illumina) for generations 1-3 were generated as previously described (9).

### STR genotyping

We used HipSTR to genotype 1,476,299 autosomal STRs, including 777,273 homopolymers (52). We ran HipSTR in its default mode, which performs *de novo* stutter estimation and STR calling with *de novo* allele generation. Since HipSTR was designed to learn locus-specific PCR stutter models, which will differ as a function of the sequencing technology, the Element and Illumina datasets were genotyped independently using the same HipSTR parameters. Output VCFs were filtered to remove calls if any of the following conditions were met: 1) a posterior < 90%, 2) > 15% of reads have a flank indel, 3) > 15% of reads have a stutter artifact, or 4) genotypes with an allele or strand bias p-value < 0.01.

### De novo mutation identification and validation

We first sought to identify candidate DNMs for each trio in the pedigree using HipSTR genotypes. On a per-trio basis, we queried each STR locus for a DNM except in the cases where one or more members of the trio had 1) a missing genotype or 2) fewer than 15 STR-spanning reads at that locus. For loci that meet these criteria, any allele observed in the child but absent in both parents is considered a candidate DNM. At loci that pass these criteria, if the child has a genotyped allele that is absent in both parents, this is considered a candidate DNM.

After genotype-based DNM discovery, we applied read-based filtering to identify high-confidence DNMs. After filtering, an average of 45.6%% of the Element candidate DNMs remain, compared to an average of only 2.6% of Illumina candidate DNMs, owing largely to a pool of candidates that is, on average, nine times larger than the Element pool. Candidate DNMs were validated by read-level inspection using a custom Python pipeline. For each candidate DNM, we extracted primary alignments from BAM files for the child and both parents using pysam (53) and computed the net difference in inserted or deleted bases within the repeat region by parsing the CIGAR string of each read. Reads were required to have a minimum mapping quality (MAPQ) of 60 and to fully span the repeat locus (±1 bp padding). For each individual, we summarized the number of reads supporting each distinct net indel size. Extensive visual inspection was used to calibrate additional thresholds. For the Element data, we required ≥2 (Q50) or 3 (Q40) reads supporting the de novo allele in the child and ≤1 read matching the de novo allele in the parents. For Illumina data, more stringent filtering was necessary. We required a minimum of 6 reads supporting the de novo allele and no evidence of the de novo allele in the parents.

### Determining parent of origin for DNMs

For a subset of individuals for whom both parents and offspring are included in the pedigree, we used a well-described technique to assign validated DNMs to a parent of origin. In brief, for each DNM in a focal individual, we identified which offspring inherited the mutant STR allele. We then scanned the surrounding genomic region of informative SNVs, which we define as a variant that is heterozygous in the focal individual, present in only one parent, absent in the focal individual’s spouse. We cataloged which children inherited the informative SNV and assessed whether inheritance of the STR DNM and informative SNV were concordant across children. If the STR allele co-segregated with the informative SNV haplotype, we inferred the parent of origin. We only assigned a parent of origin at loci where the most frequent transmission pattern occurred at ≥75% of informative SNVs.

### Regression models

We fit Poisson regression models to calculate the effect of allele length, nucleotide content, and parental age on the DNM rate. The log of the number of loci successfully tested for a DNM was used as an offset. For the allele length model, we binned homopolymer loci by their reference allele length (bp), which was used as a proxy for the allele that mutated. For the nucleotide composition model, homopolymers, dinucleotide repeats, and trinucleotide repeats were classified by the simplest version of their repeat motif. For the parental age effect model, we limited the model to trios for which DNMs could be assigned to a parent of origin. Models were fit independently for maternal and paternal ages.

## Declarations

### Ethics approval and consent to participate

Informed written consent was obtained from all individuals in this study. The study was approved by University of Utah Institutional Review Board (IRB protocol IRB_00065564). Forty-one members of the pedigree provided consent for biobanking with controlled access. Seven individuals provided consent for access to individual-level data via a Data Use Agreement with the University of Utah.

### Consent for publication

Not applicable.

### Availability of data and materials

All underlying data from 41 members of the pedigree are available with controlled access through dbGaP (phs004339.v1). Data from 7 participants (200009, 200013, 200090, 200110, 200113, 200116, 200134) are available from the corresponding author upon reasonable request and with a Data Use Agreement. Custom code and pipelines used in this study are publicly available at GitHub (https://github.com/hannah-happ/homopolymers_platinum_plus).

### Competing interests

The authors declare that they have no competing interests.

### Funding

This work was supported by the National Human Genome Research Institute (T32HG008962 to HCH, R01HG012252 to ARQ) and the Eunice Kennedy Shriver National Institute of Child Health and Human Development (R01HD106112 to ARQ).

### Authors’ contributions

HCH, DWN, and ARQ contributed to the design of the project and interpretation of results. HCH, TAS, and ARQ developed the methodology. DW prepared libraries and performed AVITI sequencing. DWN oversaw participant recruitment. HCH created the figures. HCH and ARQ wrote the manuscript with input from all authors. DWN and ARQ supervised the project.

## Acknowledgements

We thank the Utah Genome Project, the University of Utah Center for Genomic Medicine, and Element Biosciences for sequencing support. Our computational resources were partially supported by an NIH Shared Instrumentation Grant 1S10OD021644-01A1. We also thank Harriet Dashnow, Michael Goldberg, Michelle Noyes, and members of the Quinlan lab for their helpful feedback and discussion on this project. Finally, we thank the CEPH/Utah individuals for their participation.

## Supplementary Information

Additional file 1: Supplementary figures and tables. Figs. S1-S3, Table S2

Additional file 2: Supplementary table. Table S1.

## Notes

https://github.com/hannah-happ/homopolymers_platinum_plus

### Competing Interest Statement

The authors have declared no competing interest.

