## Additional file 1 for "AVITI sequencing of a four-generation CEPH/Utah pedigree confirms low mutation rates at homopolymer loci despite their low sequence complexity"

**Figure S1. Previously reported genome-wide STR mutation rates.** Mutation rates as a function of STR motif size.

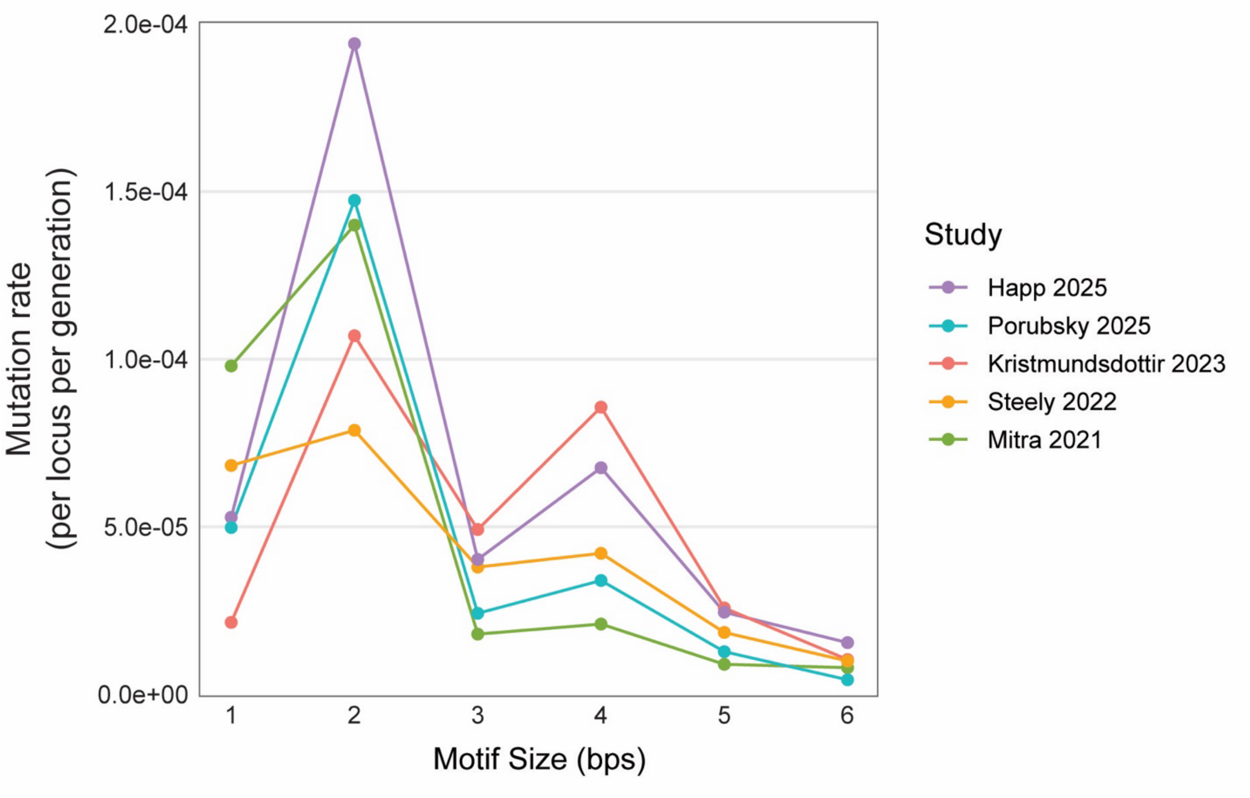

**Figure S2. Correlation of observed parental allele lengths at homopolymer loci and the allele length in the reference genome (GRCh38).**

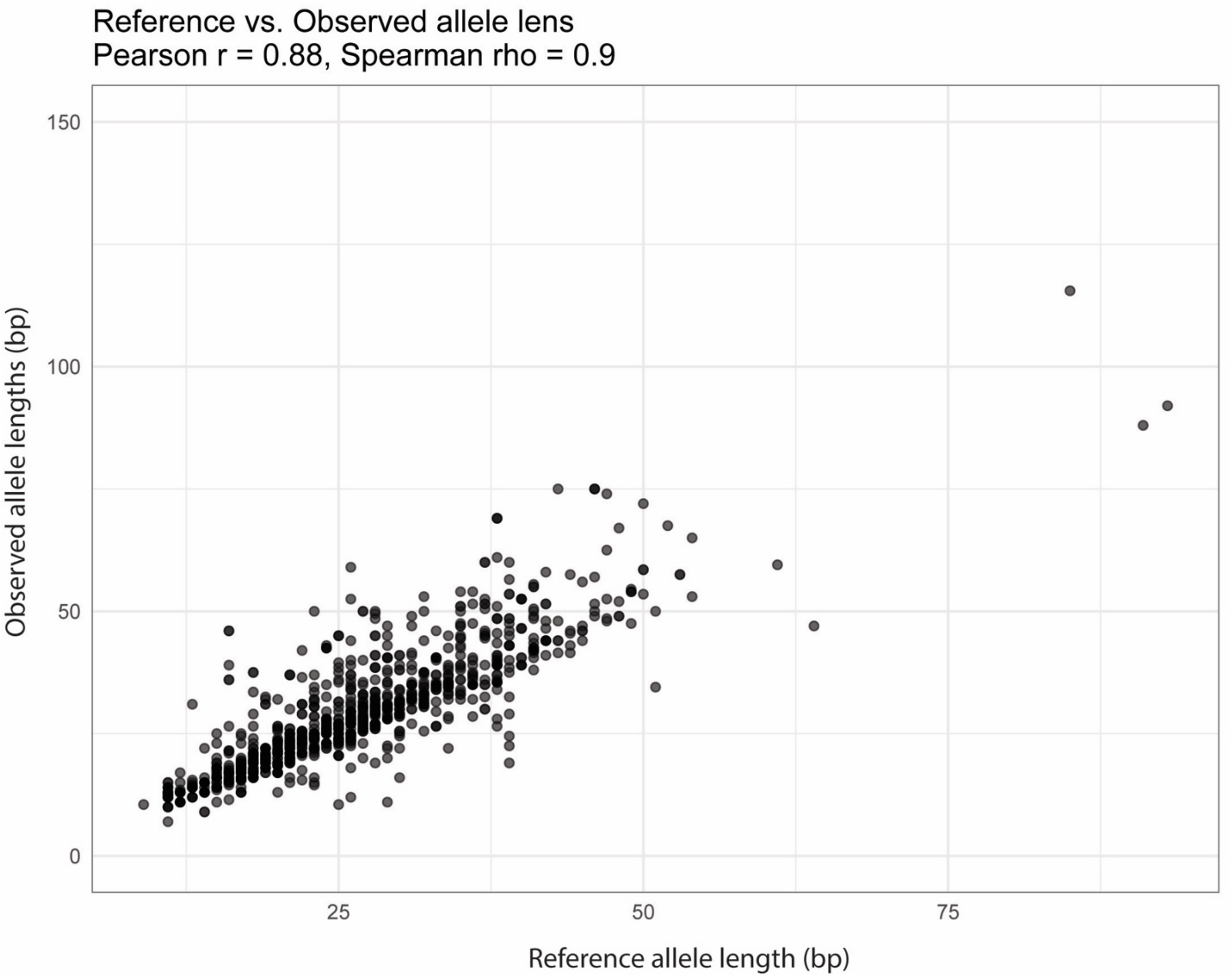

**Figure S3. Mutation rates across homopolymer, dinucleotide, and trinucleotide loci as a function of motif nucleotide content.** Homopolymers were grouped by base content (A/T, G/C), dinucleotide motifs were collapsed by reverse complement (e.g. AG = CT) and trinucleotide motifs were collapsed by shifting and reverse complementation (e.g. AAC/GTT = ACA/TGT = CAA/TTG).

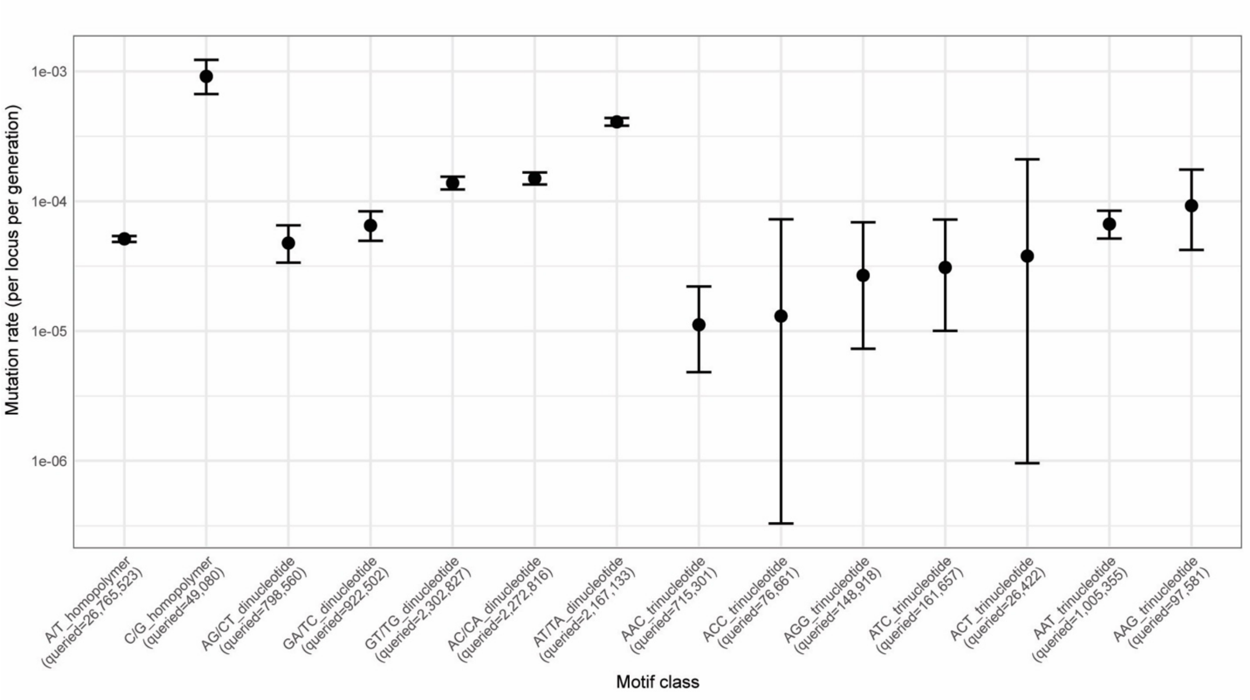

**Table S2. Homopolymer *de novo* mutation rates stratified by length bin.**

| Length bin (bp) | DNMs | Queried sites | Mutation rate (per locus per generation) | 95% CI |
|---|---|---|---|---|
| 10-24 | 727 | 23280405 | $3.12*10^{-5}$ | $2.90*10^{-5}$ - $3.36*10^{-5}$ |
| 25-39 | 594 | 3735034 | $1.59*10^{-4}$ | $1.47*0^{-4}$ - $1.72*10^{-4}$ |
| 40-54 | 90 | 379175 | $2.37*10^{-4}$ | $1.91*10^{-4}$ – $2.91*10^{-4}$ |
| 55-49 | 2 | 39669 | $5.04*10^{-5}$ | $6.11*10^{-6}$ – $1.82*10^{-4}$ |
| 70-84 | 0 | 6043 | 0 | 0 – $6.1*10^{-4}$ |
| 85-99 | 3 | 822 | $3.65*10^{-3}$ | $7.53*10^{-4}$ – $1.07*10^{-2}$ |
